# Viruses and vectors tied to honey bee colony losses

**DOI:** 10.1101/2025.05.28.656706

**Authors:** Zachary S. Lamas, Frank Rinkevich, Andrew Garavito, Allison Shaulis, Dawn Boncristiani, Elizabeth Hill, Yan Ping Chen, Jay D. Evans

**Affiliations:** USDA-ARS Bee Research Lab, BARC-East Bldg. 306, Beltsville, MD, 20705, USA; USDA-ARS Honey Bee Breeding, Genetics, and Physiology Laboratory, Baton Rouge, Louisiana, United States of America; USDA Office of the Chief Scientist, Washington D.C

## Abstract

Commercial beekeepers in the US reported severe colony losses early in 2025, as colonies were being staged for their critical role in the almond pollination season in California. Average reported losses since the preceding spring exceeded 60%, with substantial variation among operations. Many colonies were still actively collapsing in January, 2025, when pooled and individual samples were collected then screened for levels of known honey bee pathogens and parasites. Deformed wing virus strains A and B, along with Acute bee paralysis virus, were found at unusually high levels, either in pooled colony samples or in individual bees exhibiting shaking behaviors and morbidity.

Differences between these two analyses suggest that direct collections of morbid bees provide a superior diagnostic for causal viruses, a suggestion borne out by confirmation of symptoms and morbidity following isolation and new inoculations. Since these viruses are known to be vectored by parasitic *Varroa* mites, mites from collapsed colonies were in turn screened for resistance to amitraz, a critical miticide used widely by beekeepers. Miticide resistance was found in all collected *Varroa*, underscoring the urgent need for new control strategies for this parasite. While viruses are a likely end-stage cause of colony death, other stressors such as nutritional stress and agrochemicals may have also played significant roles.

## Introduction

As a managed pollinator, the honey bee, *Apis mellifera*, is an integral component of agriculture, providing key pollination services for a wide variety of crops. Honey bees are estimated to provide $US20-$30 billion in pollination services in the United States^1^, and $387B globally^2^. In the US, most honey bee colonies are moved by beekeepers at least twice per year to provide pollination services and to generate income from hive products^3^. Multiple stressors impact honey bee health throughout this annual cycle. Biotic stressors include *Varroa destructor* and other parasitic mites, viruses, and gut parasites, while pesticides present a severe abiotic threat. Coupled with erratic availability of pollen and nectar, and driven by additive and synergistic impacts, these stressors are known to impact both individual and colony health^4^.

Honey bees have evolved resilience in the face of biotic^5^ and abiotic^6^ challenges inherent to beekeeping. This resilience includes rapid worker production followed by the production of new queens and colony fission events, a key trait that allows honey bees and beekeepers to rebuild populations following losses^7^. Nevertheless, managed honey bee populations regularly experience extreme local declines, involving the near or total collapse of populations. Termed ‘disappearing disease’^8^ or ‘Colony Collapse Disorder’^9^, these sudden losses can be locally devastating. Specifically, after severe loss events surviving colonies are insufficient in number or condition to repopulate an area resulting in intense financial and emotional strain on commercial beekeepers and those dependent on pollination services^7^. Without stable populations, these losses reverberate through intertwined and codependent agricultural communities. Historically, these losses have been highest over winter months, when adult worker bees must persist for months before queens begin laying eggs and the colony is able to rebuild. More recently, summertime mortality has become a venerable component of annual losses^10-14^, pushing overall loss rates above 50% in recent years^13^.

A suite of viruses, primarily positive, single-stranded RNA viruses, predominate in managed colonies in the continental United States^15^, and are associated with both healthy and impacted colonies. A dicistrovirus, Israeli acute paralysis virus, was associated with sudden colony losses in some colonies in early 2007^16^, and additional dicistroviruses Kashmir bee virus and Acute bee paralysis virus were linked to weak colonies in a more extensive survey of this event, albeit inconsistently across regions^17^. While localized honey bee colony losses have been documented throughout the history of beekeeping, they have been especially frequent following the arrival in a region of the invasive honey bee ectoparasitic mite *Varroa destructor*^18^. This nearly ubiquitous mite vectors some, but not all, honey bee viruses, and has been a major synergist for Deformed wing virus (variants DWV-A and DWV-B), a member of the viral family *Iflaviridae*, which includes positive-sense single-stranded RNA viruses commonly infecting insects. DWV invariably increases in prevalence and magnitude after the introduction of *Varroa* to an area^19^,often followed by high colony loss rates ^20,21,22^.

In January, 2025, commercial beekeepers began reporting severe losses in commercially managed operations. These losses were reported just prior to almond bloom, the largest pollination event in the world during which more than 1.5 million colonies are staged in the Central Valley of California, USA, for a one-month flowering season^23^. Losses occurred for colonies that had spent the prior year throughout the US and that were stored both outdoors and in massive temperature-controlled ‘sheds’. As losses unfolded it was evident that over 60% of commercial beekeeping colonies had been lost since the prior summer, representing 1.7 million colonies with an estimated financial impact of $US600 million (https://www.projectapism.org/colony-loss-information).

Here we describe analyses based on six large commercial beekeeping operations which experienced these severe losses. The operations represent a collective summertime high of 183,750 managed colonies, roughly 6.8% of all managed colonies in the United States. We provide a detailed descriptive study of colony conditions, morbidities, and operation losses in light of management records and summertime locations. Quantitative analyses of parasites and pathogens point to RNA viruses as a key companion of morbid bees and commercial colonies. Critically, viruses were indicated in both pooled samples from surviving colonies and in individual bees showing behavioral morbidities. Viruses were further implicated via experimental inoculation experiments that both replicated behavioral symptoms seen in the field and caused rapid mortality of adult bees. As in past loss events^16,17^, these viruses belong to a subset of bee viruses vectored by the widespread bee parasitic mite, *Varroa destructor*.

Additional analyses of *Varroa* mites showed the universal presence of a genetic marker for resistance to the widely used acaricide amitraz^24^ in mites from collapsed colonies, highlighting the difficulties inherent in controlling mite populations. While pesticides and other stressors are likely to be additional factors involved in bee declines, these results suggest that actions to mitigate the impacts of parasitic mites and their associated viruses are critical.

## Materials and Methods

### Field inspections and collections

Samples were collected as described previously^7^. In brief, all samples were collected individually when not collected as pooled samples, and directly onto dry ice. Symptomatic bees dying in front of colony entrances were collected before internal colony inspections. Colonies were given a sample identification number and were scored for strength and observable morbidities. A random sample of adult bees from the center of the brood nest was collected as a pooled sample for parasite, pathogen and pesticide analysis. Bee brood (larvae and pupae) was visually inspected for signs of morbidity and both healthy and symptomatic individuals were collected along with any attached *Varroa* mites. Bee bread (stored pollen-based food), wax, honey, and ‘entombed pollen^25^ were also recorded and collected.

### RNA Extraction and cDNA Synthesis for Pathogens

RNA was extracted from both pooled samples and individual bees. A pooled sample of fifty bees from each colony was used for molecular detection of common bee parasites and pathogens using established protocols^26^ Briefly, each pooled sample was placed in a disposable RNA extraction bag (Bioreba, Reinach, Switzerland) and 25 mL aliquot of guanidine thiocyanate lysis buffer was added prior to manual homogenization with a marble rolling pin on a flat board. After homogenization, total RNA was extracted using an acid phenol protocol for bulk bees. After extraction, 5 µL aliquots of total RNA were treated with DNase I (Invitrogen, Waltham, MA) at 37°C for 1 hour followed by 75°C for 10 minutes, as per manufacturers’ instructions. First-strand complimentary DNA (cDNA) was generated using iScript™ Advanced cDNA Synthesis Kit, following manufacturers protocol. The synthesized cDNA was diluted 1:5 with molecular grade water to provide a template in qPCR reactions.

Individual bees were extracted using a Trizol-chloroform precipitate extraction. A 1 mL aliquot of Trizol was added to each individual bee in a screw cap microcentrifuge tube with 10ul of 1mm sterilized silica beads, and agitated in a bead mill homogenizer at 4000 RPM twice for 30 seconds with one minute between agitation cycles to prevent samples from heating. A 200ul aliquot of chloroform was added to each microcentrifuge tube and then shaken by hand twice for 15 seconds. Samples were allowed to incubate for two minutes at room temperature, and then were centrifuged at 4°C and 12,000 RPM for 20 minutes. The supernatant was transferred to 1.7ml Eppendorf tubes containing 1ml of molecular grade isopropyl and then incubated at -20°C overnight. Eppendorf tubes were vortexed briefly and centrifuged as previously described.

Isopropyl was removed, then pellets were washed with 80% molecular grade ethanol with 15 minutes centrifuge cycles during each wash. Pellets were air dried and then dissolved in 50ul of molecular grade water.

### Pathogen Quantification

qPCR was performed using SsoAdvanced™ Universal SYBR® Green Supermix (Bio-Rad, Hercules, CA) in a CFX96™ Real-Time System thermal cycler (Bio-Rad, Hercules, CA) following all manufacturers’ instructions. The cycling parameters are as follows: initial denaturing at 95°C for 30 seconds, followed by 50 cycles of 95°C for 5 seconds and 60°C for 30 seconds. A melt curve was generated for each sample following the completion of the run protocol for amplicon validation. Supplementary Table S1 describes PCR primers for all screened targets and controls. Absolute quantification of target copy number was calculated by creating a standard curve using standards with known concentrations (provided by the National Honey bee Health Survey^27^), calculating the slope, dilutions and then calculating concentration in samples (*Supplementary Table S2 and S3*).

### Inoculum preparation and infectivity

Viral inoculum was prepared from individual field collected specimens. Symptomatic specimens, exhibiting erratic behavior, immobilization or inability to fly outside of colony entrances were collected individually and saved on dry ice until permanent storage at -80°C. Viral inoculum was prepared by separating viral particles from host material through repeated freeze-thaw cycles.

Samples were processed individually by grinding with a sterile pestle in 1% phosphate buffered solution, followed by freezing at -80°C. Each sample underwent a total of three freeze-thaw cycles, concluded by bringing the total volume of PBS to 1500ul. Finally, samples were pressed through a 0.22um filter and then stored at -80°C for long term storage. Total RNA was extracted from a 50ul aliquot of inoculum using an RNEASY kit to manufacturer’s instructions. 5ul of RNA was used as a template for cDNA preparation using a one-step iScript kit (Biorad), incorporating DNASE1 and RNASEOut to manufacturer’s instructions.

Inoculum infectivity was tested by direct injection into purple-eye pupae procured from an overwintered colony at USDA-ARS BRL managed apiaries in Beltsville, MD. Inoculum was diluted 10-fold in PBS and then 1ul of diluted inoculum was delivered to each pupae using a Word Precision microinjector. Total delivery was made via a 8ul injection. Pupae were incubated at 32°C and 60% RH. Pupae were collected over three time series: time of injection (Time Zero), 36 hours post-injection and 60 hours post-injection then were flash frozen at -80°C until total RNA extraction.

### Longevity impacts of collected inocula

Inocula were tested for virulence on adult bees in controlled cage trials. A frame of newly emerging bees was collected from a single colony at USDA-ARS BRL managed apiaries in Beltsville, MD, and incubated at 32°C and 60% RH. Bees were allowed to emerge overnight, and then were pooled and randomized between treatment groups. The study included a PBS control injection, and four inoculum experimental injections. Additionally, an aliquot of each experimental inoculum was heat-inactivated at 95C for one hour, serving as an additional control. Each inoculum consisted of three concentrations, resulting in a total of 17 groups (12 inocula, 4 heat inactivated, 1 PBS). 16 bees were injected per group for a total of 272 individual adult bees. A no-inject negative control was utilized (N = 16) to indicate if damage was incurred by the shunt injection. Adult bees were housed in P-cups ^28^ with a fondant feeder and incubated at 32°C and 60% RH. Bees were inspected two hours post-injection, and then every twelve hours thereafter for a total of 15 days. Morbidity and mortality were recorded at each inspection. An additional trial, lasting seven days, was carried out to test survivorship of a single, virulent inoculum (CV5) using concentrations 10^−7^-10^−9^ bee equivalents. This trial also utilized a heat inactivated control, PBS and negative control (N = 172).

### Statistical analyses

We used baseR and RStudio with the accompanying packages: Vegan, MASS, DescTools, lmerTest, survival, survminer and GGPlot for statistical analysis and figure creation. Our data from pooled diagnostic screening and individual bee diagnostic screening did not meet the conditions for normality. As a result, non-parametric test statistics were utilized. Due to the nature of a case study, descriptive states were used when sample collection was limited or when an experimental group was not available. A Permutational Multivariate Analysis of Variance (PERMANOVA) using Euclidean distance and 999 permutations was used to measure group means and variances for pooled bee samples across the whole dataset, and across operations using the condition of the colony as a predictor variable. A non-parametric Mann-Whitney U test (Wilcoxon rank-sum test) was utilized to measure if there was a significant difference in pathogens detected between strong and weak colonies within an operation. Pathogen abundance and prevalence was calculated using descriptive statistics, and then we analyzed the relationship between the abundance of detected pathogens and pathogen load for each colony to calculate the Spearman rank correlation coefficient. For analysis on individual bee specimens, we used the Mann-Whitney U test to measure significant differences of viral load between symptomatic (dying) and asymptomatic bees. Analysis was done using morbidity as a sole grouping variable for individual bee analysis. A Linear mixed-effects model using Satterthwaite’s method for approximating degrees of freedom for t-tests was fitted for each virus (DWV-A and DWV-B) separately to investigate if source (beekeeper operation) was significantly associated with either virus, using operation as the fixed effect and individual specimens as a random intercept to account for potential non-independence of samples within colonies. Two-way cluster analyses for viral titers in healthy and morbid bees were carried out via hierarchical clustering (SAS-JMP version 18, Cary, NC).

Infectivity tests for viral inocula were analyzed on the group level using a one-way analysis of variance. Survivorship assays were analyzed using a log-rank test to compare survivorship amongst treatments and presented using a Kaplan-Meier survivor analysis.

### Screening for genes linked to amitraz resistance in honey bees

Individual mites were assayed for a genetic marker linked to amitraz resistance^24^. 16 mites were collected from five recently collapsed colonies while 23 mites were collected from live- colony brood cells containing developing bee larvae. Mites were stored at -80°C until shipping to the USDA-ARS Honey Bee Breeding Genetics and Physiology Lab in Baton Rouge where they were again stored at -80°C until processing. Individual *Varroa* mites were transferred into 2 mL collection microtubes (Qiagen 19560) with a single sterilized 5 mm steel bead and 60 µL of nuclease free water. Samples were then homogenized during 4 cycles on a TissueLyser II (Qiagen) at 30cycles/s for 10s followed by 5 second intervals with rotation of the tubes between cycles. After homogenization, they were centrifuged for 6 minutes at 2272xg. The supernatant was used immediately for genotypic determination as below.

Allelic discrimination of susceptible (Y215) and resistant alleles (Y215H) of the β2 octopamine receptor (Octβ2R) was done utilizing TaqMan technology according to previously published methods ^29^. The region of the Octβ2R containing the amitraz resistance mutation was amplified using forward (5′-GGA TAC CGT GCT CAG TAA TGCT-3′) and reverse (5′-CTG TCG GGT CGC TTC TAG ATAG-3′) primers (standard oligonucleotides with no modification). With the VIC® labeled fluorescent probe (5′-ATG CGC CAA TAA GTG AAT -3′) for the detection of the wild-type allele, and the 6FAM™ labeled probe (5′-CGC CAA TGA GTG AAT- 3′) for detection of the Y215H mutation ^29^. Each probe also had a 3′ non-fluorescent quencher and a minor groove binder at the 3’ end.

TaqMan assays were assessed using a Bio-Rad CFX Connect with 10µL reaction volumes comprised of 5 uL of TaqMan mastermix (ThermoFisher), 0.5 uL of Taqman assay which includes the labeled probes and primers, and 4.5 uL of DNA extract. Samples were held at 95°C for 10min followed by 40 cycles of 95°C 10s then 60°C for 30s. Wild type and mutant strains were confirmed with wells containing plasmid DNA. Genotypes were determined using CFX Maestro (BioRad) and exported to an Excel spreadsheet for analysis.

## Results

### Pathogen and parasite levels

We examined pathogen loads in pooled samples from 72 weak colonies and 41 strong colonies. The median number of pathogen detections was 5 (IQR = [2]) for both weak (n = 72) and strong colonies (n = 41). A total of 532 pathogenic targets were detected (Table 1). The most prevalent virus in these samples was Deformed wing virus (78%), a longstanding bee pathogen that has surged worldwide thanks to the vectoring of parasitic mites. The dicistrovirus Acute bee paralysis virus, also mite-vectored, showed unusually high levels compared to prior surveys (e.g.,^27^), and was found in 72% of screened colonies. Surprisingly, while presence of viruses was notably high in these affected operations, viral load did not differ between pooled samples from weak and strong colonies (Figure 1).

**Table 1.** Prevalence of key honey bee pathogens and parasites across sampled colonies. Focal pathogens are highlighted by bold font.

| Pathogen | Detection (n = 113 colonies) | Prevalence |
| --- | --- | --- |
| **Acute bee paralysis virus (ABPV)** | **81** | **71.68%** |
| Black queen cell virus (BQCV) | 85 | 75.22% |
| Chronic bee paralysis virus (CBPV) | 5 | 4.42% |
| **Deformed wing virus-A (DWV-A)** | **88** | **77.88%** |
| **Deformed wing virus-B (DWV-B)** | **19** | **16.81%** |
| Israeli acute bee paralysis virus (IAPV) | 5 | 4.42% |
| Kashmir bee virus (KBV) | 32 | 28.32% |
| Lake Sinai virus (LSV) | 72 | 63.72% |
| Nosema cerana (NOS) | 71 | 62.83% |
| Sac brood virus (SBV) | 40 | 35.40% |
| Trypanosome (TRYPS) | 34 | 30.09% |

**Figure 1:**
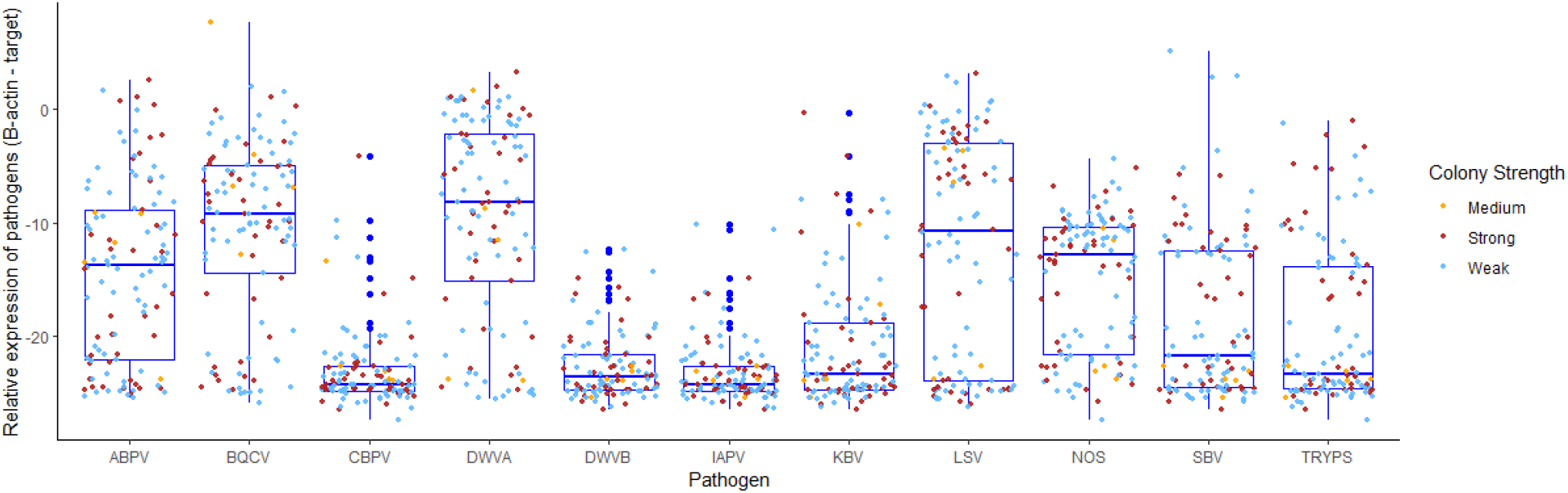
Diagnostic screening of pooled adult bees from surviving colonies (N = 113). There was no significant difference in pathogen loads between strong and weak colonies across all colonies in the study (Permanova, F(2,110)=0.79,R2=0.014,p=.663). There was no significant difference in individual pathogen loads within operations as well (Mann-Whitney U test result with Bonferroni correction).

To better assess virus connections to real-time morbidity we screened symptomatic and asymptomatic bees from individual colonies. Virus levels were strikingly higher in bees showing signs of behavioral impairment indicative of disease or pesticide exposure. In total, 38 symptomatic adult bees were paired with 28 asymptomatic (control) bees. Individual dying bees had significantly higher levels of DWV-A and DWV-B, W=254, p = .0006, W=11, p < .0001 (Figure 2). DWV-B was not detected in symptomatic controls, but was detected in 100% of symptomatic adults (*Supplementary Table S4)*.

**Figure 2:**
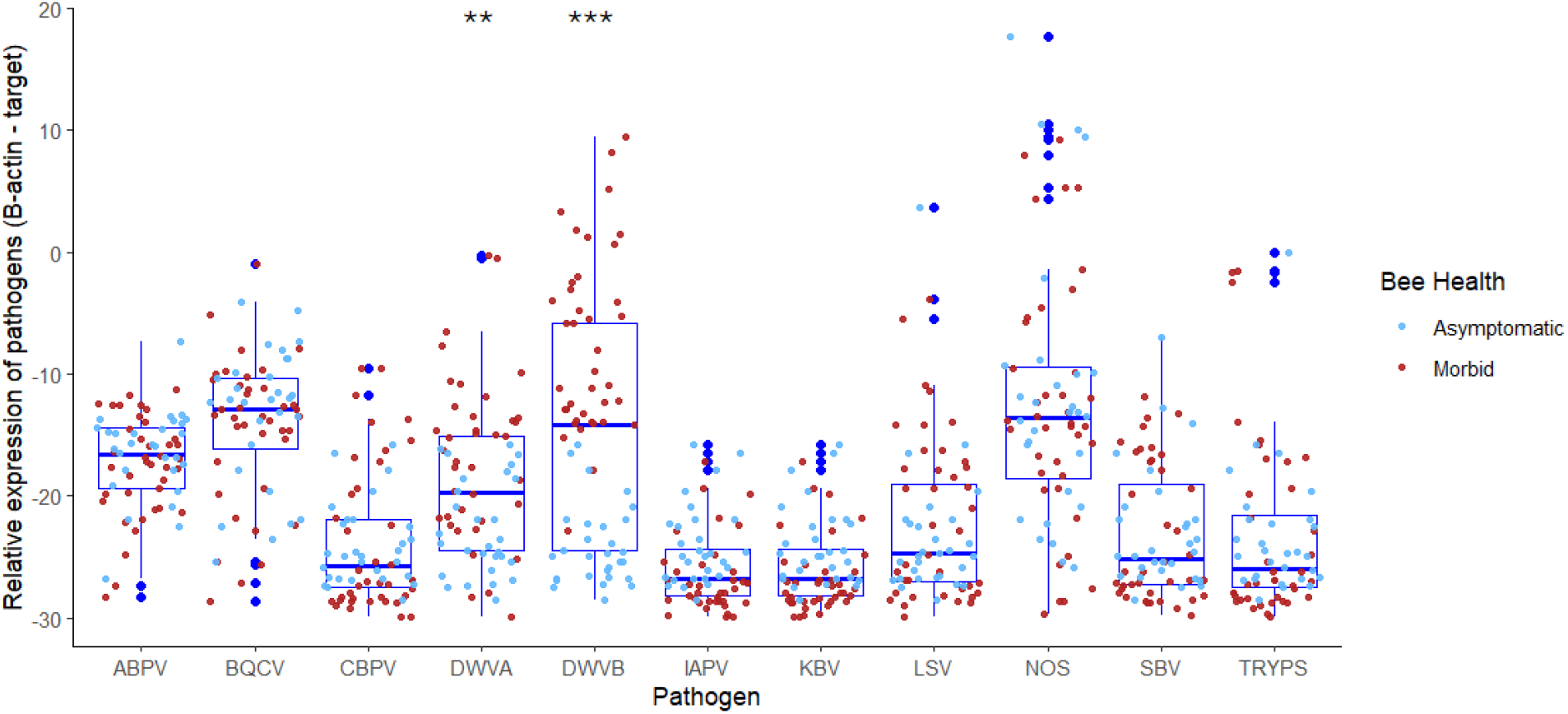
Diagnostic screening of individual adult bees, showing significantly higher levels of DWV-B in symptomatic individuals.

The source of individual bees (beekeeper operation) did not have a statistically significant effect on viral targets DWV-A or DWV-B. A linear mixed-effects model revealed no statistically significant effect of by operation (F(2, 4.308) = 1.918, p = 0.257 for the viral target DWV-A in morbid samples (n=38). Pairwise comparisons showed no significant differences in DWV-A between Operation 2 and Operation 3 (estimate = 4.311, SE = 4.403, t(2.029) = 0.979, p = 0.430) or between Operation 5 and Operation 3 (estimate = 7.396, SE = 4.127, t(4.292) = 1.792, p = 0.143). A direct comparison of Operation 3 also showed no significant difference between Operation 5 and Operation 2 (estimate = 3.085, SE = 4.979, t(2.337) = 0.620, p = 0.591). No statistical significant effects were found, respectfully, for the viral target DWV-B: Operation (F(2, 4.346) = 1.369, p = 0.343, nor through the same respective pairwise comparisons (estimate = 9.297, SE = 4.902, t(2.343) = 1.897, p = 0.179) or (estimate = 7.633, SE = 4.426, t(4.286) = 1.725, p = 0.155) or (estimate = -1.664, SE = 5.500, t(2.605) = -0.302, p = 0.785). A two-way hiearchical clustering analysis (Figure 3) shows that correlative viral patterns differed across morbid bees. While DWV was generally linked to morbidity, and both DWV strains clustered together, some cohorts of morbid bees had high levels of both viruses while others had high levels of only DWV-B.

**Figure 3:**
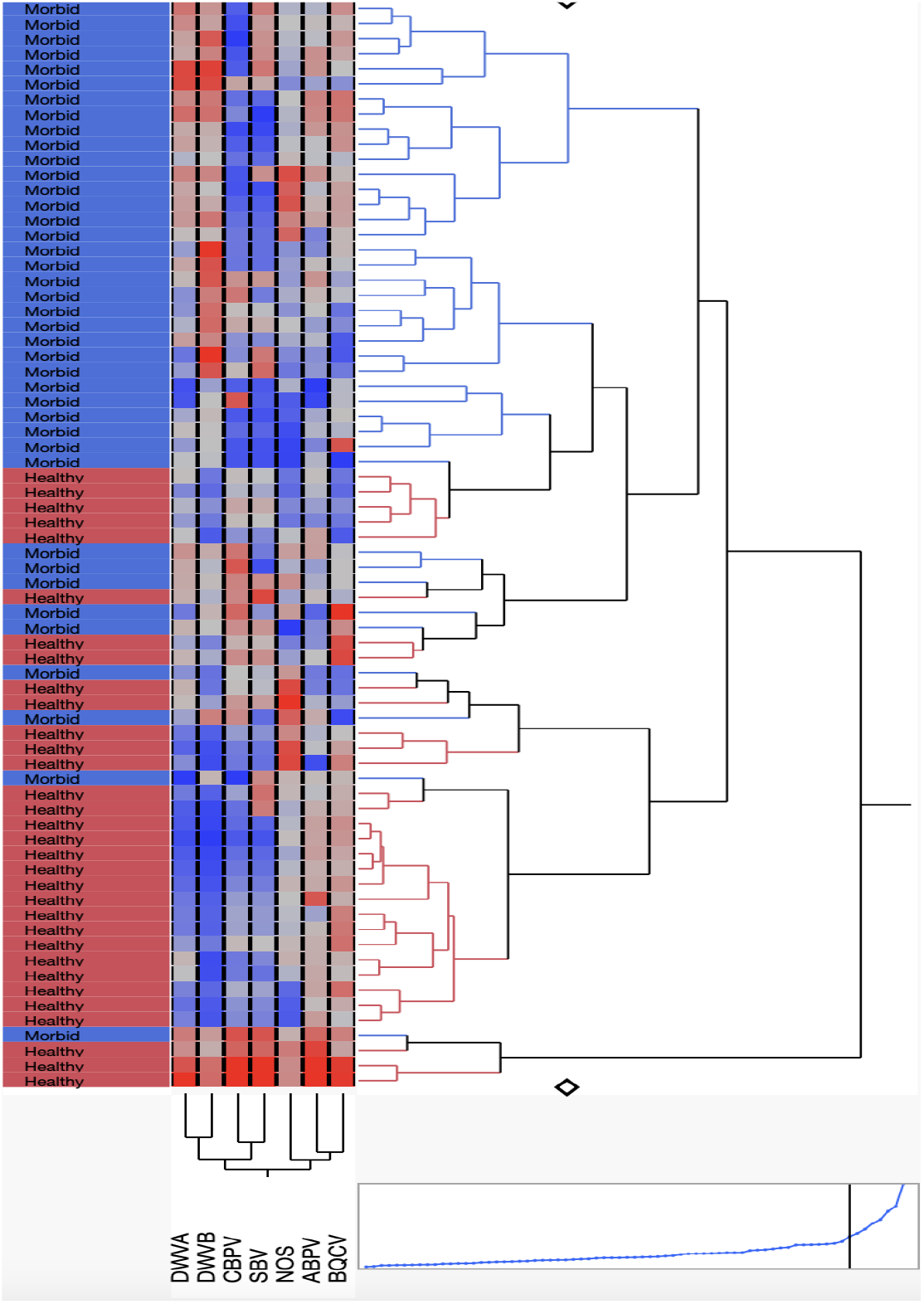
Two-way cluster analysis showing levels (Blue low to red high) of seven pathogens found in individual bees exhibiting healthy or morbid behaviors.

**Figure 3.**
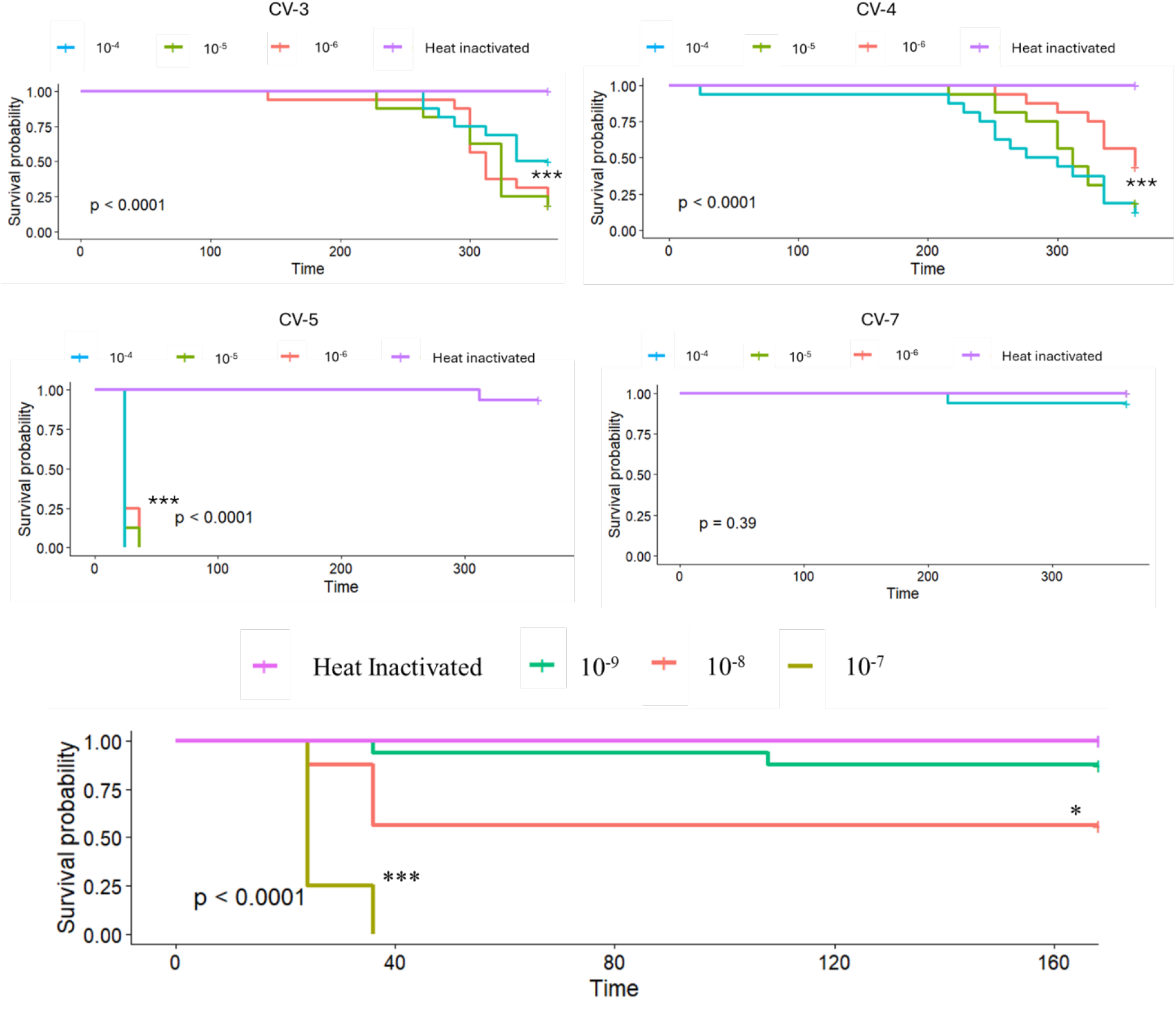
a) Time-course survival for naive bees inoculated with four inoculum sourced from field collected individual honey bees, presented at concentrations of 1.5 × 10^−4^ to 10^−6^ bee- equivalents b) Survival of bees injected with serial dilutions of inoculum CV5, from initial concentration to concentrations of 1.5 × 10^−7^, 10^−8^, and 10^−9^ of the original isolate.

**Figure 4.**
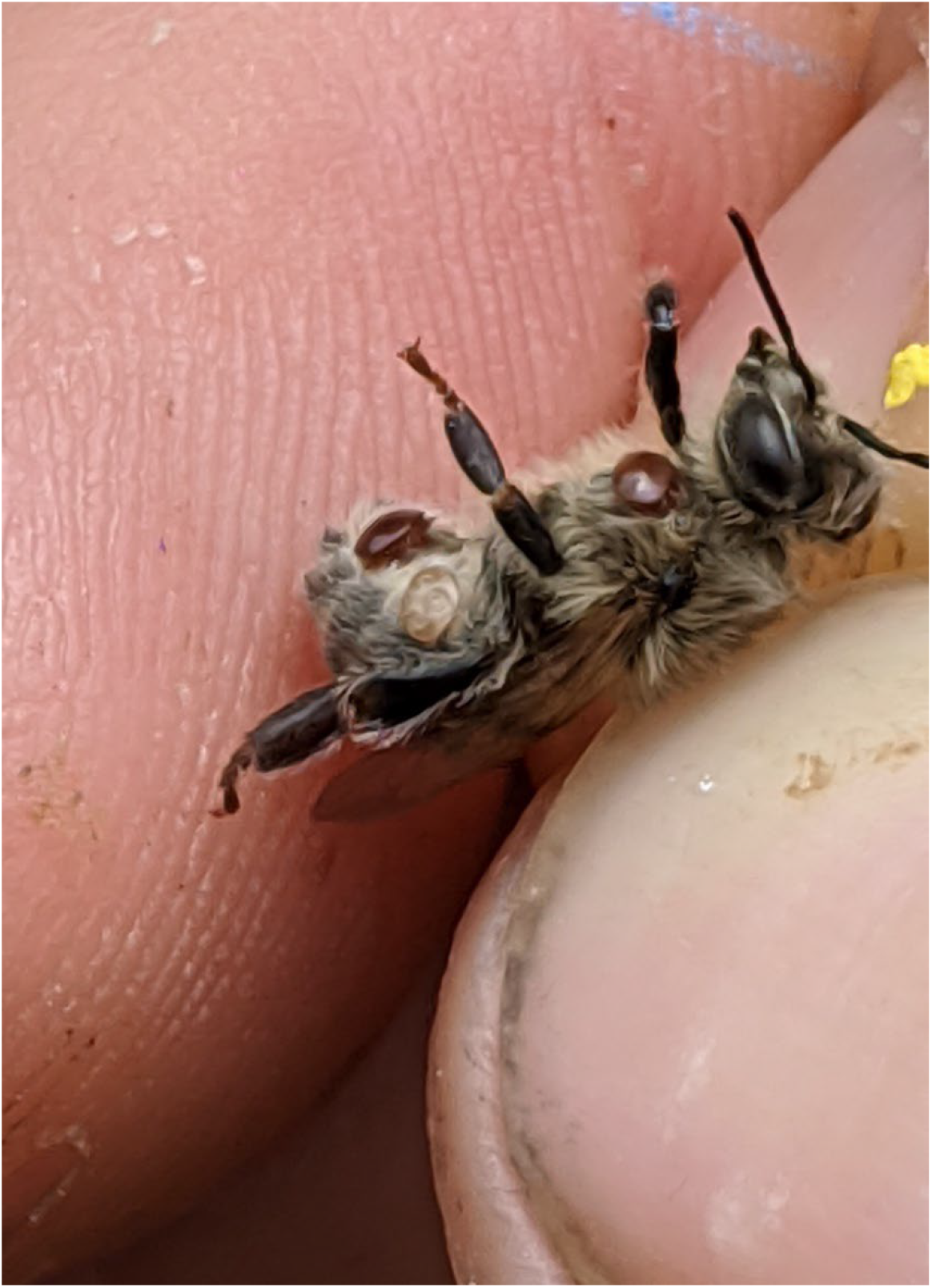
A heavily parasitized emergent adult honey bee showing females mites that were parasitic during larval and pupal development.

### Infectivity and Induced mortality from viral inocula

Isolates purified from individual bees showing behavioral morbidities (CV3, CV4, CV5 contained a range of known bee viruses (*Supplementary Table S3)*, with isolate CV5 showing especially high levels of ABPV (3.52 × 10^7^ GE/ul). Comparatively the other two did not have detectable levels of ABPV, but instead had high levels of DWV-B (CV3 = 8.43 × 10^6^ GE/ul, CV4 =2.27 × 10^6^ GE/ul) An isolate purified from an asymptomatic bee showing no behavioral morbidities did not have detectable levels of ABPV, DWV-A or DWV-B (CV7 = No detection). All three isolates sourced from morbid bees caused infectivity in injected pupae (*Supplementary Figure S1*) at 36 and 60 hours post-injection (ANOVA, Table 2A), and significantly higher than heat-inactivated controls (ANOVA, Table 2B). DWV-B was amplified in each inoculum sourced from morbid bees, while ABPV was also robustly amplified in pupae injected with CV5. CV7, the isolate sourced from a single asymptomatic bee, had amplification of the DWV-B target. However, levels at 36 and 60 hours post-injection were not significantly higher than samples immediately frozen at injection (Time-Zero, T_0_), indicating that viral amplification did not significantly increase over the delivered dose.

**Table 2:** Infectivity of four inoculum 36 and 60 hours post-injection compared to injections at T_0_ and comparatively to viral levels of heat inactivated controls.

| **Table 2A: Anova table, inoculum infectivity compared to Time-Zero injections** |  |  |  |  |  |  |
| --- | --- | --- | --- | --- | --- | --- |
| **Inoculum** | **Viral Target** | **Sum of Squares** | **Df** | **Mean Square** | **F** | **P** |
| CV3 | DWV-B | 940.8 | 1 | 940.8 | 240.7 | <0.0001 |
| CV4 | DWV-B | 1072.4 | 1 | 1072.4 | 505.5 | <0.0004 |
| CV5 | DWV-B | 280.27 | 1 | 280.27 | 39.26 | 0.0002 |
| CV5 | ABPV | 1795 | 1 | 1795 | 565.1 | <0.0001 |
| CV7 | DWV-B | 100.9 | 1 | 100.87 | 1.987 | 0.201 |
| **Table 2B: Anova table, inoculum infectivity compared to heat inactivated controls** |  |  |  |  |  |  |
| **Inoculum** | **Viral Target** | **Sum of Squares** | **Df** | **Mean Square** | **F** | **P** |
| CV3 | DWV-B | 1846 | 1 | 1845.9 | 25.01 | 0.0001 |
| CV4 | DWV-B | 1696 | 1 | 1696 | 20.93 | 0.0004 |
| CV5 | DWV-B | 495.7 | 1 | 495.7 | 17.35 | 0.0006 |
| CV5 | ABPV | 2143 | 1 | 2143.2 | 19.95 | 0.0003 |
| CV7 | DWV-B | 577.5 | 1 | 577.5 | 13.2 | 0.0022 |

The single isolate from an asymptomatic bee neither showed amplification of viral targets in pupae, nor caused significant loss in adult bees. However, the three isolates sourced from morbid bees all had negative impacts on survivorship when injected into naive bees at a concentration of 10^−4^ - 10^−6^ bee-equivalents (Figure 3, *Supplementary Table S5*). Isolate CV5 was especially virulent, inducing 100% mortality by 36 hours in every dilution factor in Experiment 1 (10^−4^ to 10^−6^ bee-equivalents). An additional trial was run, showing that this isolate significantly affected bee survivorship inducing 100% mortality in every trial when diluted to a factor of 10^−7^ bee- equivalents. When diluted to 10^−8^ bee-equivalents CV5 induced 44% mortality, suggesting that, when injected, this isolate would represent an approximate LD_50_ dose with enough infectious material from a single sourced bee capable of killing ca. 66 million adult bees.

### *Amitraz resistance in* Varroa *mites*

In total, 39 individual mites, retrieved from 18 colonies in five beekeeping operations, were screened for a robust marker for amitraz resistance. Mites were collected from surviving (N = 13) and perished colonies (N= 5). Of these, 100% had resistance genotypes, a trait that was fard less common in the most recent widespread survey^24^.

## Discussion

Here we show a preponderance of RNA viruses in bees collected in the midst of a colony collapse event that impacted a majority of US managed honey bee populations. High levels were found for two iflaviruses known to cause honey bee mortality^30^ and an unusually high incidence was recorded for the dicistrovirus Acute bee paralysis virus. These three viruses are transmitted between bees via the parasitic mite *Varroa destructor*. This mite is ubiquitous in bee colonies and has risen to high levels in recent surveys. One widely used chemical control for *Varroa*, amitraz, is suspected of losing efficacy after decades of heavy use^24^. In analyzing mites from collapsed colonies, all screened mites showed a genetic marker associated with amitraz resistance. All beekeepers in our screening relied upon amitraz for the majority of their miticide applications, underscoring the challenges faced by beekeepers in controlling mite damage.

Controlled laboratory assays showed that viral inocula procured from dying bees led to replicative viruses and high pathogenicity. The iflaviruses DWV-A and DWV-B were present in three of these four inocula. One inoculum, labeled CV5 and derived from a single bee showing behavioral signs of disease, had relatively high levels of Acute bee paralysis virus alongside DWV-A and DWV-B. This inoculum was especially deadly when injected into naïve bees, killing exposed bees even when diluted 10^−8^-fold from its original suspension, resulting in a 44% mortality rate for exposed bees. By extrapolation, the suspension from this single bee was sufficiently pathogenic to cause the mortality within days of ca. 66 million exposed bees. To find no effect, a 10^−9^-fold dilution was made, resulting in no significant difference in mortality when compared to PBS or heat inactivated controls. The fact that a tenfold difference in dilution lead to such a drastic change in mortality (44% to 0%) highlights the extreme virulence of this viral combination and the steepness of its dose-response curve.

*Varroa destructor* is considered the most impactful factor for honey bee colony survivorship ^18^, especially when coupled with damaging RNA viruses. *Varroa* affects honey bees on the individual, colony and population levels. Through their direct feedings on brood and adult bees, *Varroa* efficiently vector viruses directly into individuals^31-33^. *Varroa* engage in a rapid feeding behavior as they actively switch amongst adult worker bees in order to feed^34^. This has been shown to cause intense pressure on the worker bee cohort, where a small number of mites are capable of parasitizing most members of a populous honey bee colony^35^. Without viruses, these feedings impart little hazard directly to the parasitized bee. However, *Varroa* efficiently vector a suite of honey bee viruses, which subsequently transmit through numerous horizontal routes host-to-host^36^. Viral infected bees live shorter lives. When the premature death of adult bees surpasses the rate of replacement through brood rearing, colonies risk collapsing. Crashing colonies pose risk to neighboring colonies as they become a source for *Varroa* and viral dispersal in densely packed operations^37,38^.

*Varroa* parasitism has been kept in check largely by beekeeper management and the prolific use of combative miticides^39^. Adequate *Varroa* control is necessary for honey bee colony survival as *Varroa* are one of the most significant factors associated with high rates of overwinter colony losses around the world ^40-44^. Coumaphous and tau-fluvalinate were two effective miticides which fell out of use when mites developed resistance. Beekeepers have largely been reliant upon amitraz since then due to its low impact on bee health, low cost and ease of use. Unfortunately, amitraz resistance in *Varroa* is now widespread^45^ and the Y215H mutation in Octβ2R, which is the putative receptor for amitraz, is consistently associated with amitraz resistance^46^. The universal prevalence of this resistance genotype in all of these samples is strongly suggestive that amitraz may have been ineffective at controlling these *Varroa* populations. The removal of amitraz as a viable tool to manage *Varroa* will destabilize beekeeping operations until a replacement miticide or management strategy arrives.

Sudden losses of worker bees from established honey bee colonies have been noted for decades. Described as ‘disappearing disease^8^ or, more recently ‘Colony Collapse Disorder’^9^, these events have often remained unsolved, with populations gradually rebuilding in subsequent years^47^. In early 2007, widespread colony losses in the United States were heavily scrutinized for biotic causes, with multiple viruses and gut parasites emerging as correlates with weakened colonies. One dicistrovirus, Israeli acute paralysis virus, was linked with colony losses in early samples from this loss episode^16^, while additional viruses were predominant in weak colonies in more extensive surveys^17^,^9^.

A cross-sectional study design has limitations, in that prior exposures and specimens lost before sampling result in knowledge gaps. This fact can be partially overcome by carrying out longitudinal studies such that potential causes are recognized prior to becoming existential threats to colonies ^21^. Given the suddenness of these losses, we were not able to collect samples prior to most losses. Instead, colonies were sampled directly as they emerged from indoor overwintering sheds or from outside areas with reduced reproduction and activity prior to the full spring. More importantly, we focused on individual bees that were expressing behaviors known to precede death by minutes or hours. This sampling regime highlights the ‘survivorship bias’ inherent to sampling loss events and arguably helps clarify the challenges faced in pinpointing causes for prior colony loss events^16,17^. Coupled with our infection bioassays, we are confident that these results point to a causal factor for a large fraction of honey bee colony losses.

Honey bees suffer from a combination of biotic and abiotic stresses and we cannot rule out the importance of additional causes in these declines. Efforts are ongoing to generate and analyze unbiased RNA and DNA sequencing resources for these bees to scout for known and novel parasites and pathogens. Pesticides are known to exacerbate the impacts of *Varroa* and other disease agents in reducing colony success^48,49^ and it is entirely possible that such synergisms have compounded the risk to honey bee colonies. Accordingly, bees and hive substrates are being screened for signs of known agrochemicals. Nevertheless, this study describes a surprisingly high prevalence of bee viruses, including ABPV, and a direct link between viral loads and native or induced morbidity. These active infections in surviving colonies, coupled with the seasonal increase in *Varroa* levels, present continued risk this year for managed colonies. Arguably, this supports a near-term shift in mite treatment strategies, while also minimizing additional stressors. A systems approach to protecting and managing honey bees from all threats is needed in order to maintain honey bees as a key player for worldwide agriculture.

