## Supplementary Information for "Viruses and vectors tied to honey bee colony losses"

### Supplementary Tables and Figures

**Table S1.** Primer names and associated sequences for qPCR detection and relative quantification of common honey bee pathogens.

| Target | Primer | Sequence |
|---|---|---|
| Actin | AMActin.F | TTGTATGCCAACACTGTCCTTT |
| Actin | AMActin.R | TGGCGCGATGATCTTAATTT |
| Deformed Wing Virus A | DWV.F | GAGATTGAAGCGCATGAACA |
| Deformed Wing Virus A | DWV.R | TGAATTCAGTGTCGCCCATA |
| Deformed Wing Virus B | VDV1.F | GCCCTGTTCAAGAACATG |
| Deformed Wing Virus B | VDV1.R | CTTTTCTAATTCAACTTCACC |
| Nosema ceranae | qNC40sRP.F | AGAAACTACAACAGCATCACTGGGA |
| Nosema ceranae | qNC40sRP.R | AGTGAATATTCCAATTCCCAACGACTT |
| Acute Bee Paralysis Virus | ABPV.F | ACCGACAAAGGGTATGATGC |
| Acute Bee Paralysis Virus | ABPV.R | CTTGAGTTTGCGGTGTTCCT |
| Chronic Bee Paralysis Virus | CBPV.F | CAAAATCAACGAGCCAATCA |
| Chronic Bee Paralysis Virus | CBPV.R | AGTGTGAGGATCACCGGAAC |
| Sacbrood Virus | SBV.F | GGGTCGAGTGGTACTGGAAA |
| Sacbrood Virus | SBV.R | ACACAACACTCGTGGGTGAC |
| Israeli Acute Paralysis Virus | IAPV.F1a | GCGGAGAATATAAGGCTCAG |
| Israeli Acute Paralysis Virus | IAPV.R1 | CTTGCAAGATAAGAAAGGGGG |
| Kashmir Bee Virus | KBV.F | TGAACGTCGACCTATTGAAAAA |
| Kashmir Bee Virus | KBV.R | TCGATTTTCCATCAAATGAGC |
| Trypanosomatidae | Trypan1.F | CTGAGCTCGCCTTAGGACAC |
| Trypanosomatidae | Trypan1.R | GTGCAGTTCCGGAGTCTTGT |
| Tracheal Mite | AcwdCO1.F | TCAATTTCAGCCTTTTATTCAAGA |
| Tracheal Mite | AcwdCO1.R | AAAACATAATGAAAATGAGCTACAACA |
| Lake Sinai Virus | LSVrsc.F | GTCATCCCAAGAGAACCACTYAC |
| Lake Sinai Virus | LSVrsc.R | CRCACYGACATGAAGAAATGAGGTC |
| Vitellogenin | VgMC.F | AGTTCCGACCGACGACGA |
| Vitellogenin | VgMC.R | TTCCCTCCCACGGAGTCC |
| Hymenoptaecin | Hymenopt.F | CTCTTCTGTGCCGTTGCATA |
| Hymenoptaecin | Hymenopt.R | GCGTCTCCTGTCATTCCATT |

| **Table S2.** Primer names and associated sequences for qPCR detection and absolute quantification of four pathogens in experimental inoculum. | | |
|---|---|---|
| **Target** | **Primer** | **Sequence** |
| Deformed Wing Virus A | DWV-F8668 | TTCATTAAAGCCACCTGGAACATC |
| Deformed Wing Virus A | DWV-B8757 | TGGCGCGATGATCTTAATTT |
| Deformed Wing Virus B (VDV-1) | qVDV1-CP-F | CTGTAGTTAAGCGGTTATTAGAA |
| Deformed Wing Virus B (VDV-1) | qVDV1-CP-R | GGTGCTTCTGGAATAGCGGAA |
| Black Queen Cell Virus | BQCV-qF7893 | AGTGGCGGAGATGTATGC |
| Black Queen Cell Virus | BQCV-qB8150 | GGAGGTGAAGTGGCTATATC |
| Acute Bee Paralysis Virus | ABPV-F6548 | TCATACCTGCCGATCAAG |
| Acute Bee Paralysis Virus | KIABPV-B6707 | CTGAATAATACTGTGCGTATC |

**Table S2:** Primer sequences used for quantification of viral inoculum.

| **Table S3.** Dilution factor of inoculum (representing portion of pupa) and copy number per ul of inoculum | | | | | | | |
|---|---|---|---|---|---|---|---|
| **Inoculum** | **Virus** | **$10^{-4}$** | **$10^{-5}$** | **$10^{-6}$** | **$10^{-7}$** | **$10^{-8}$** | **$10^{-9}$** |
| CV-3 | DWVB | 843,000 | 84,300 | 8,430 | Na | Na | Na |
| CV-4 | DWVB | 227,664 | 22,766 | 2,276 | Na | Na | Na |
| CV-5 | DWVA | 70,900 | 7,090 | 709 | 70.9 | 7.09 | < 1 |
| CV-5 | DWVB | 893,000 | 89,300 | 8,930 | 893 | 89.3 | 8.9 |
| CV-5 | ABPV | 3,520,000 | 352,000 | 35,200 | 3,520 | 352 | 35.2 |
| CV-7 | NaN | NaN | NaN | NaN | NaN | NaN | NaN |

**Table S3:** Bee equivalents and copy number per ul of inoculum.

| Table S4 Prevalence of pathogens in individually collected bees | | | |
|---|---|---|---|
| **Bee Health** | **Pathogen** | **Positive Detections** | **Prevalence** |
| **Asymptomatic (N = 28)** | ABPV | 21 | 75% |
| | BQCV | 21 | 75% |
| | DWV-A | 8 | 28.6% |
| | LSV | 1 | 3.6% |
| | NOS | 19 | 67.9% |
| | SBV | 4 | 14.3% |
| | TRYPS | 1 | 3.6% |
| **Morbid (N = 38)** | ABPV | 34 | 89.5% |
| | BQCV | 29 | 76.3% |
| | CBPV | 9 | 23.7% |
| | DWV-A | 30 | 78.9% |
| | DWV-B | 38 | 100% |
| | LSV | 16 | 42.1% |
| | NOS | 32 | 84.2% |
| | SBV | 11 | 28.9% |
| | TRYPS | 8 | 21.1% |

**Table S4:** Prevalence of pathogens found in asymptomatic and symptomatic bees.

| Table S5 Inoculums and survivorship of adult bees | | | |
|---|---|---|---|
| Inoculum | Concentration (Bee Equivalent) | Kaplan-Meier survival analysis (p – value) | Number (Percent Perished) |
| CV3 | $10^{-4}$ | $p = 0.0025$ | 8 **(50%)** |
| CV3 | $10^{-5}$ | $p < 0.0001$ | 13 **(81.2%)** |
| CV3 | $10^{-6}$ | $p < 0.0001$ | 13 **(81.2%)** |
| CV4 | $10^{-4}$ | $p < 0.0001$ | 14 **(87.5%)** |
| CV4 | $10^{-5}$ | $p < 0.0001$ | 13 **(81.2%)** |
| CV4 | $10^{-6}$ | $p = 0.0009$ | 9 **(56.2%)** |
| CV5 | $10^{-4}$ | $p < 0.0001$ | 16 **(100%)** |
| CV5 | $10^{-5}$ | $p < 0.0001$ | 16 **(100%)** |
| CV5 | $10^{-6}$ | $p < 0.0001$ | 16 **(100%)** |
| CV5 | $10^{-7}$ | $p < 0.0001$ | 16 **(100%)** |
| CV5 | $10^{-8}$ | $p = 0.005$ | 7 **(43.8%)** |
| CV5 | $10^{-9}$ | $p = 0.1673$ | 2 **(12.5%)** |
| CV7 | $10^{-4}$ | $p = 0.0009$ | 1 **(6.25%)** |
| CV7 | $10^{-5}$ | $p = 1.00$ | 0 **(0 %)** |
| CV7 | $10^{-6}$ | $p = 1.00$ | 0 **(0 %)** |

**Table S5:** Kaplan-Meier P-Values for inoculum dilutions in adult bee survivorship trials.

**Figure S1**

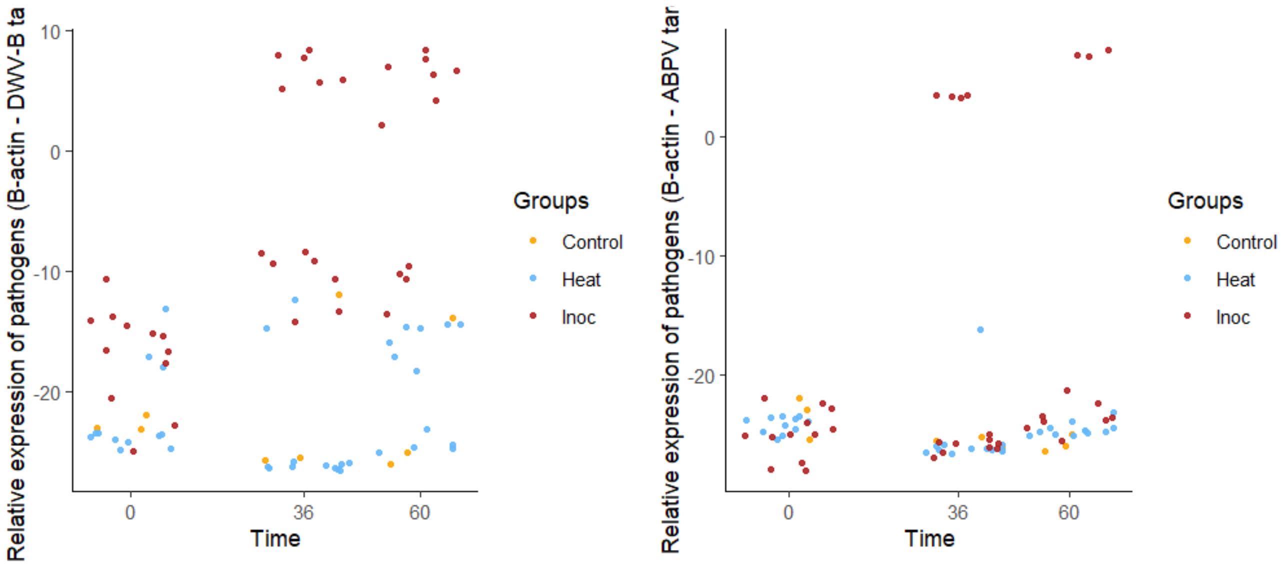


**Figure S1** Amplification of DWV-B and ABPV targets in injected pupae at time zero, 36 and 60 hours post injection between

### Colony Descriptions

Colonies generally lost the abundance of adult bees normally present within their winter cluster. The measure of these bees is not fully known due to the cross sectional design of this study. We relied upon beekeeper interviews and records to derive qualitative descriptions of loss. Below we highlight colony descriptions which convey the severity of loss while accurately portraying field observations made between January 24th and February 1st, 2025.

#### 1. Typical dwindled colony

Colonies which dwindled lost the abundance of adult bees from their winter clusters. Operation 1 colonies dwindled over the course of months. Operation 1 moved their bees from summer locations in late October to their wintering yards in California; only moving colonies in surplus of 10 frames of adult bee coverage to the state. These colonies typically were small, with a low density of adult bees covering comb surfaces. In some instances brood was not sufficiently covered by adult bees. Brood was occasionally symptomatic, though it was unclear if morbidities were from an underlying pathology, lack of attendance from insufficient worker bees, or a combination of both. Operation 1 fed sucrose to colonies in

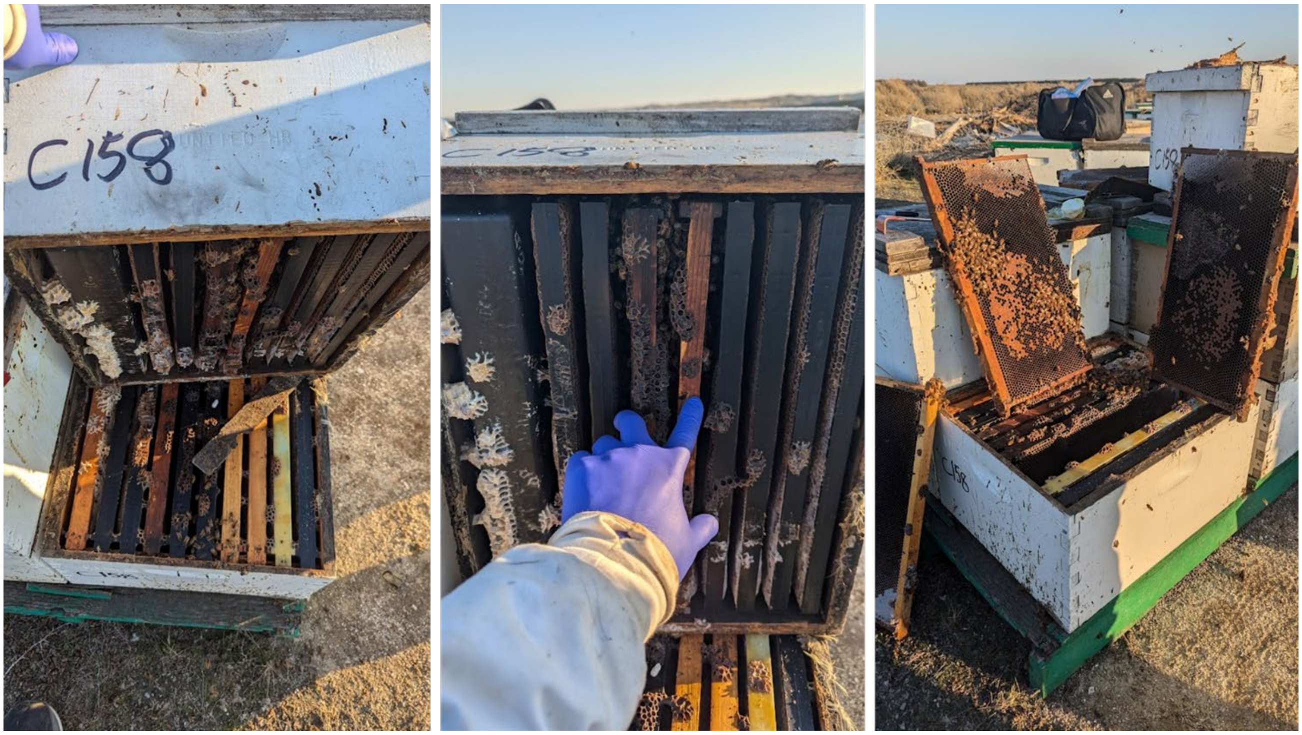


**Picture 1:** A typical ‘dwindled’ colony: honey stores largely intact, queen present, a patch of brood, and adult bee abundance lacking.

2. **Typical strong colony**

Colonies identified as strong typically had robust adult bee populations, completely covering the entire surface of the interior frames of the colony. Although quantitative estimates of adult bees were not possible, a qualitative description of dense bee coverage would accurately describe colonies rated as strong. We used strong as a qualifier during field assessments. This description is interchangeable with “unaffected” in that the colonies appeared to not suffer a rapid loss of adult bees. Brood was not a good indicator of colony strength as colonies recently removed from indoor winter storage had little, and queens in field colonies may have had intermittent laying, likely from the ebbs and flows of winter weather.

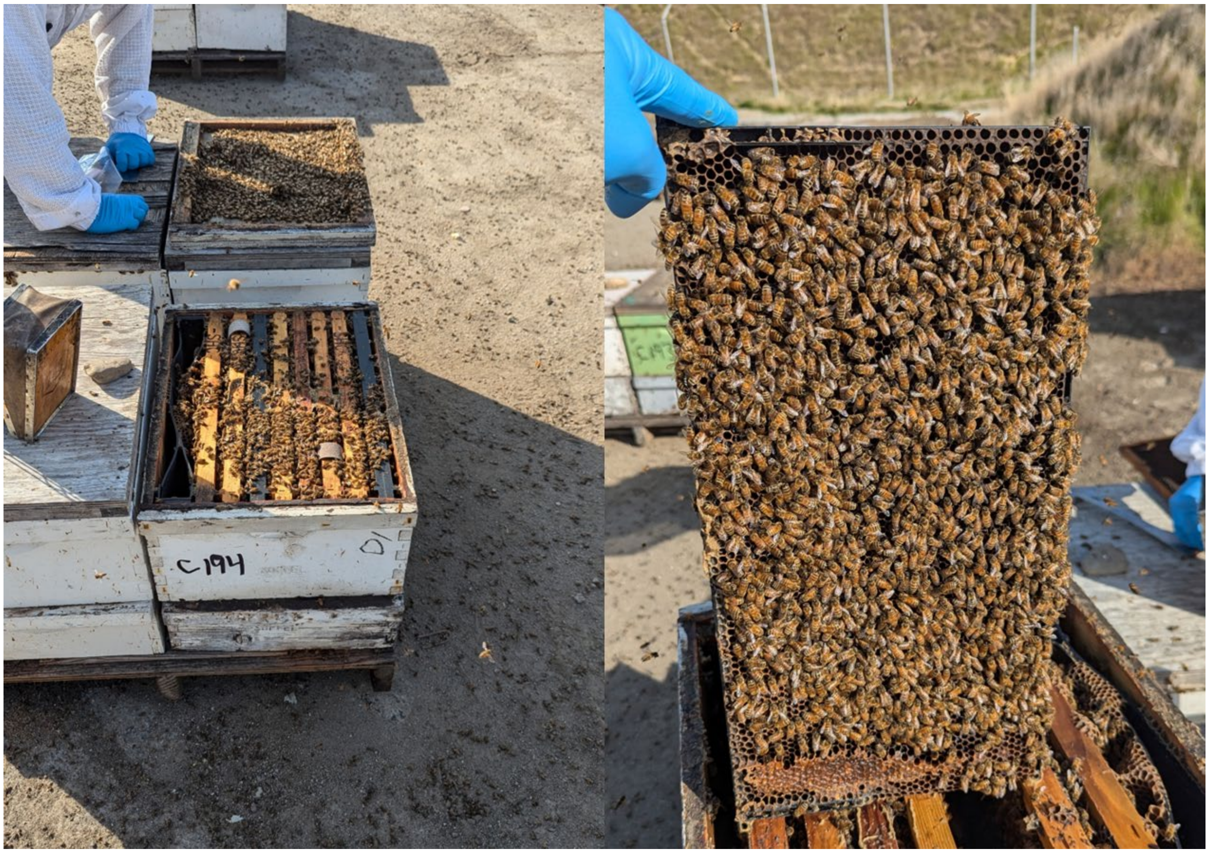


**Picture 2:** A typically strong colony, showing coverage across all internal frames, and dense coverage of adult bees on hive surfaces.

#### 3. United colonies that continue to dwindle

Beekeepers will combine similarly weak colonies with the expectation that the new unite will perform better than their component parts. We sampled united colonies in two of the six operations in this study. The success of those colonies varied widely. In Operation 3 unites faired poorly with 15 out of 18 united colonies losing bees between the time of their unification and our inspection (83.3%). In Operation 4 a similar result was observed with 46.2% of united colonies evidently losing bees (6/13). Deductions on bee loss were largely qualitative and made by deducting the duration between beekeeper unification of colonies, and the condition of the colony at our inspection.

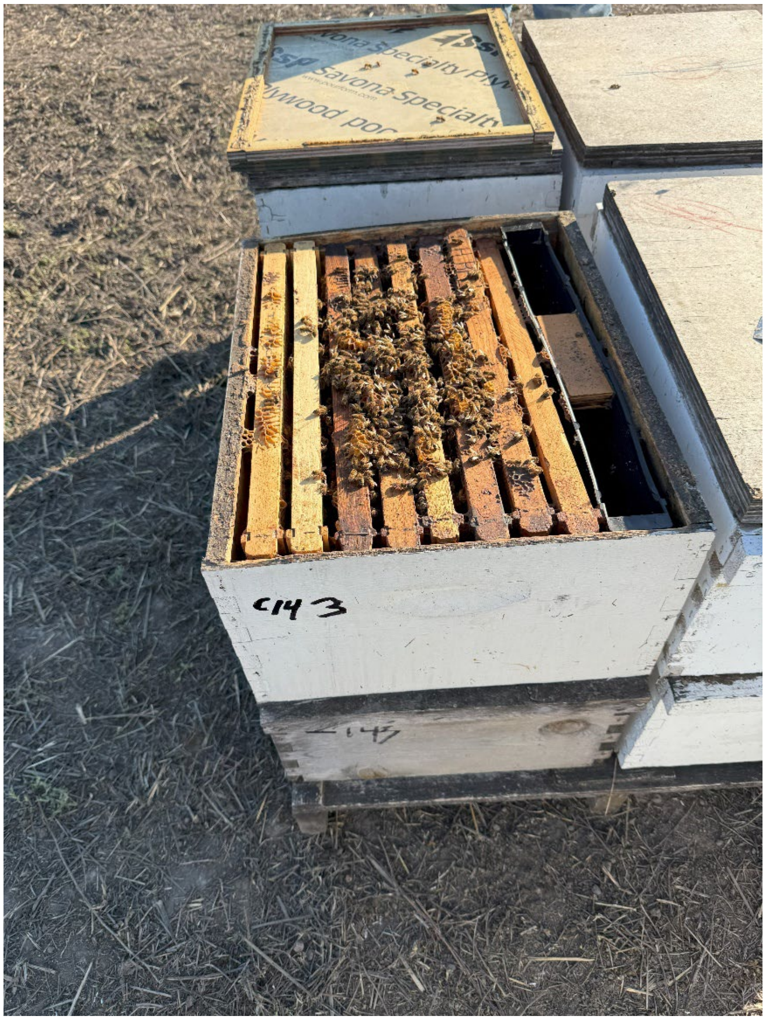


**Picture 3:**: A colony which continued to dwindle after uniting with another colony.

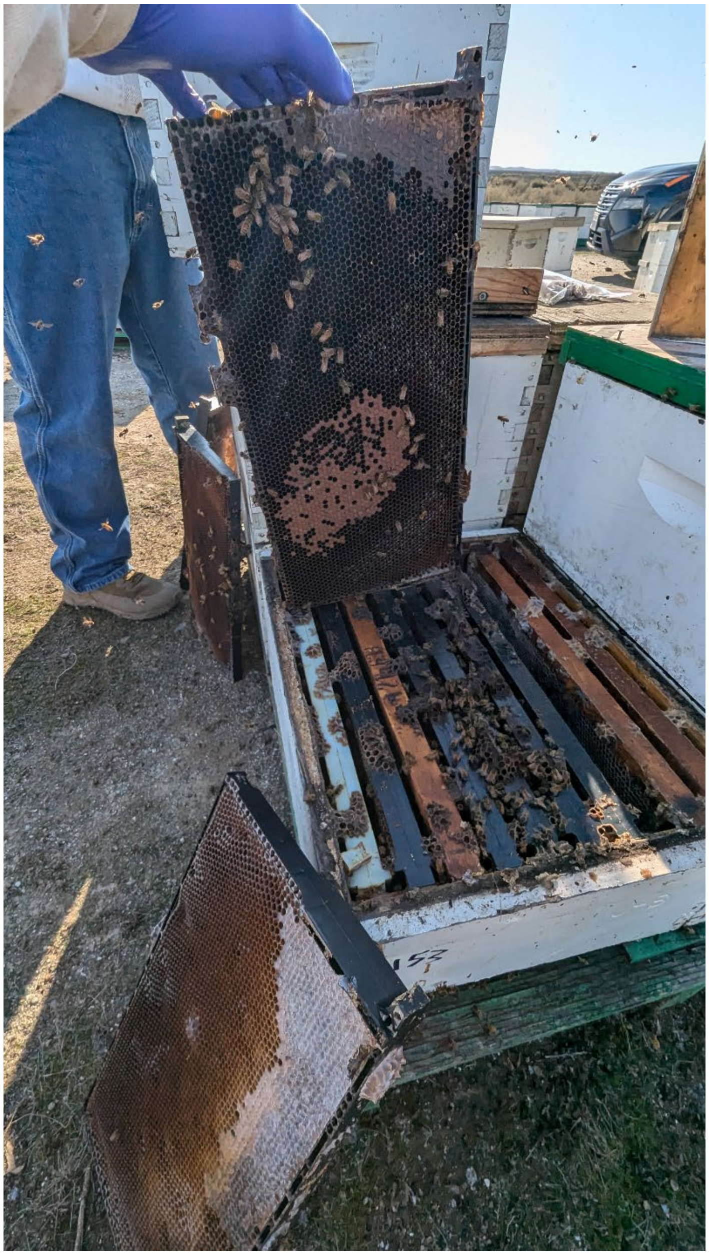

**Picture 4:** A typically “dwindled” unite is picture below. This colony lost approximately 6 frames of adult bee coverage between January 21st, 2025 when it was inspected by the beekeeper, and January 28th, 2025 when inspected by our research team.

### 4. Survivorship bias

Operation 5 overwintered colonies in climate controlled storage. Colonies were transported to CA from storage on Wednesday January 29$^{th}$, and were sampled on the morning of January 31$^{st}$. The beekeeper removed dead colonies, but left the dead bees from those colonies, including undisturbed bottom board debris. As a typical description, surviving and perished colonies had hundreds to thousands of recently dead bees exuded in the front of colonies. Dwindled colonies were typically small, with a low density of adult bees covering several frames. Brood was lacking or represented by a small patch, though that description is typical for colonies immediately out of indoor storage. Strong colonies were typically robust, with dense adult bee coverage. Brood was typical of colonies recently released from indoor storage. Perished colonies had *Varroa* visible on bottom boards and/or in the mass of adult bees outside entrances. Figure xxx is a typical representation. Every pallet bottom board inspected for debris had recoverable *Varroa*, except for two (5/7). The number of *Varroa* varied, but were never quantified. Several colonies visibly had hundreds of dead mites, while others had fewer (Figure xxx). We contrasted this qualitative observation between perished and surviving colonies as surviving colonies had few observable Varroa while remnants of perished colonies, when inspected, had many.

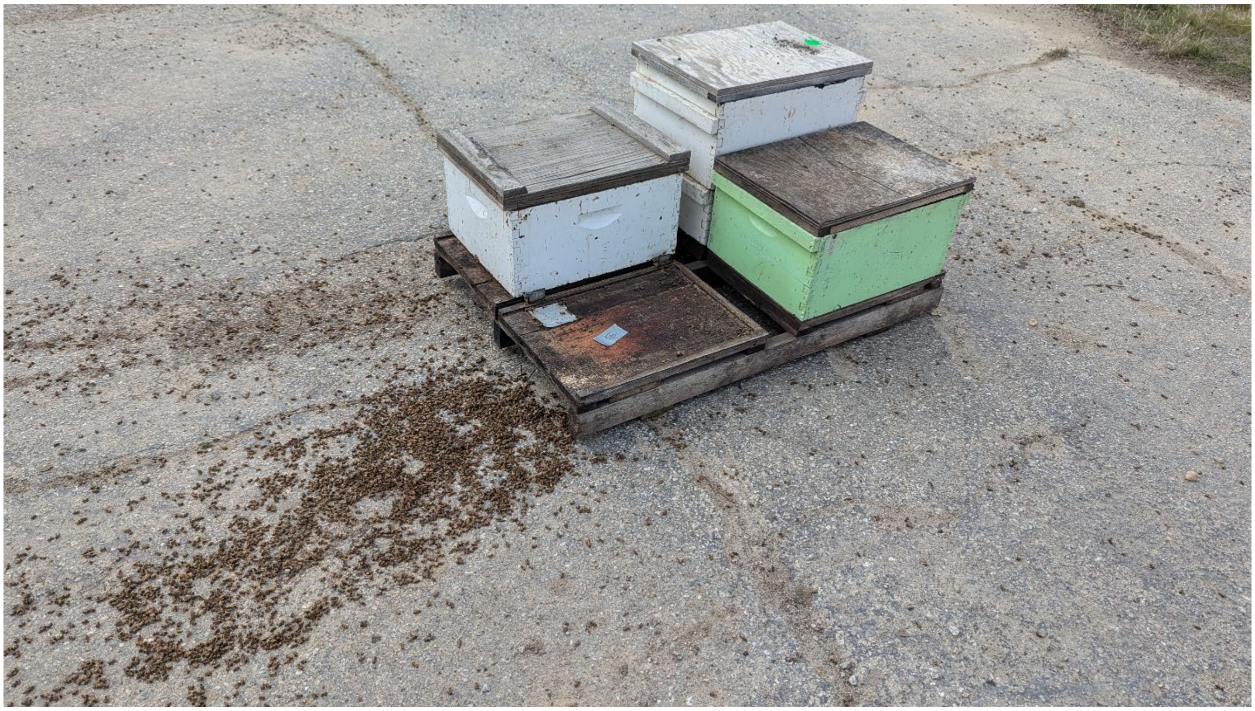

**Picture 5:**: Remains of a perished colony includes a mass of recently dead bees and pallet board debris.

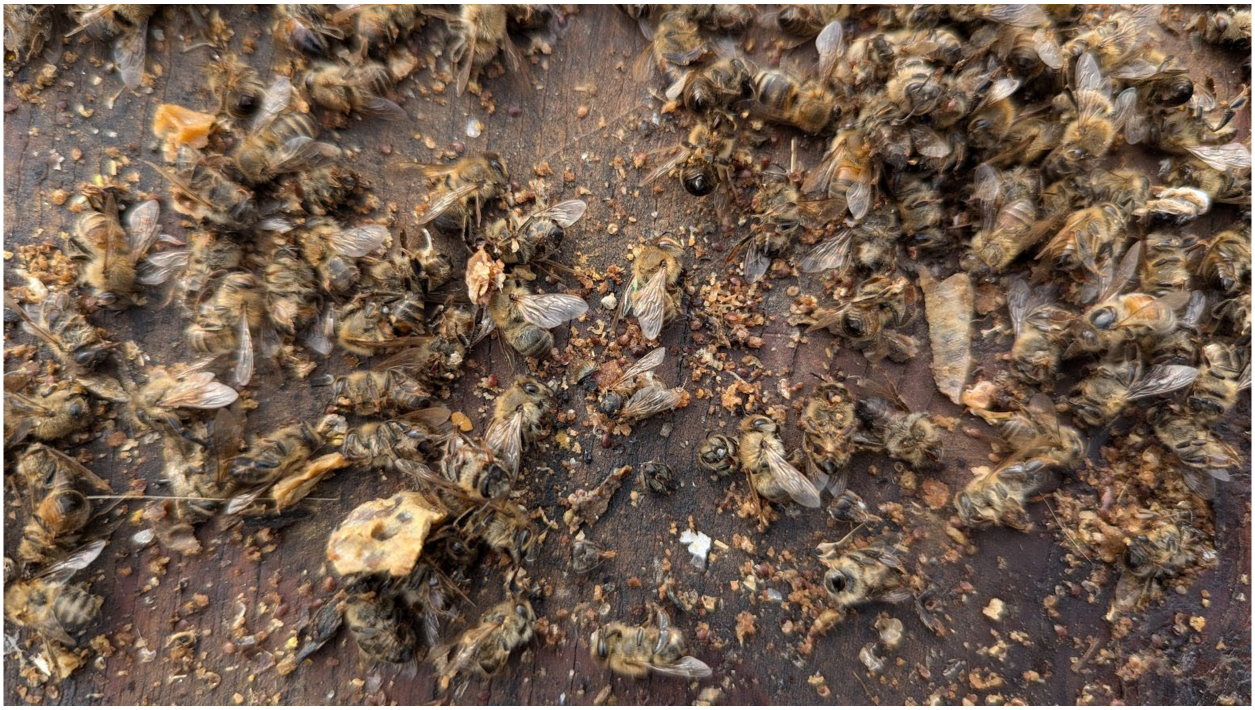

**Picture 6**: An upclose picture of pallet board debris, featuring numerous *Varroa*.

### 5. Symptomatic bees

Symptomatic bees were observed leaving from colony entrances. They were unable to fly, and instead crawled on the ground. Some became fully immobilized, appearing dead but we reactivate when warmed in the hand. Others would be able to make a short, downward flight from the colony entrance, and then crawl away from the colony. Commonly, bees would quake or shake in place or as they mobilized away from the colony. Some would climb vegetation (if present) and stay adhered to the stems. Some bees engaged in erratic movements after egressing the colony, without seemingly the ability to mobilize in a forward motion.

The rate of bees leaving colonies was unknown. In Operation 1 a carpet of degraded, perished bees, variable in size, was present in front of colony entrances. However, it is unclear if the mass of bodies accrued over a short or long duration. At Operation 2 bees could be observed leaving affected colonies every minute. Stooping over the colony entrance, one could watch morbid bees intermittently egressing. Sometimes an undertaker bee would carry out a symptomatic bee. In Operation 2 foragers, with corbicula filled with fresh pollen, were observed to lose flight and land around their colonies, unable to finish flying into the entrance.

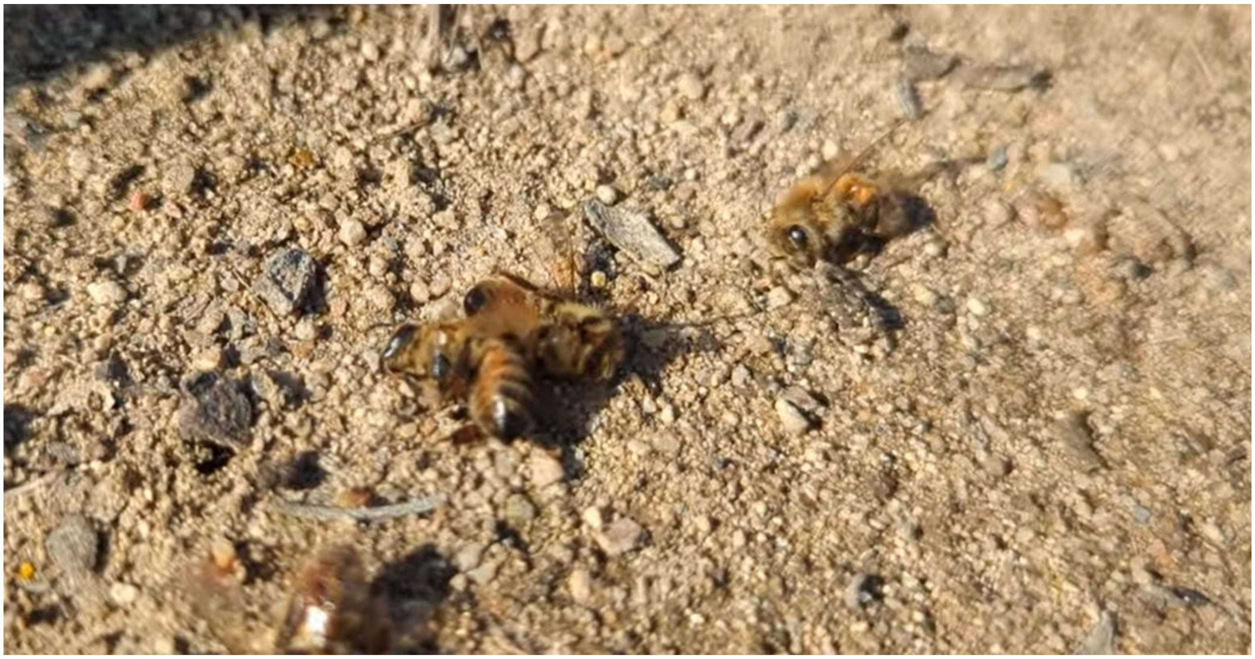

**Picture 7:** Immobilized adult worker bees, having recently egressed from their colony.
